# The role of toxin/antidote genes in the maintenance and evolution of accessory chromosomes in *Fusarium*

**DOI:** 10.1101/2025.02.05.636631

**Authors:** Linnea Sandell, Adrian Forsythe, Anna Mirandola, Samuel Jorayev, Andrew Urquhart, Alexandra Granger Farbos, Sven J. Saupe, Aaron A. Vogan

## Abstract

The genomic diversity of many fungal species is augmented by accessory chromosomes, which are variably present in individual strains. These genomic regions evolve rapidly, accumulating genes important in pathogenicity but also harboring significant amounts of transposable elements (TEs). This duality suggests a trade-off: accessory chromosomes provide infection-related benefits while otherwise being deleterious due to their highly repetitive nature and contributions to genomic instability. Despite this, accessory chromosomes often appear to be stably maintained even when strains are grown on media, with no plant host. Previously, we had observed that genes homologous to meiotic drive toxin/antidote proteins from *Podospora anserina* (*Spoks*) are abundant on accessory chromosomes in various *Fusarium* species. Using a functionality screen in yeast, we demonstrate that some of these homologs have active toxin and antidote properties. We propose that these selfish genes act to maintain accessory chromosomes during vegetative growth and may influence their spread via parasexual cycles. Finally, as *Spok* genes are mobilized by the newly described TE superfamily *Starships*, it suggests these TEs play crucial roles in forming accessory chromosomes and regions. These results illuminate a mysterious facet of fungal biology, a key step towards describing the origin, spread, and maintenance of pathogenicity in many fungal species.

## Introduction

Fungi are among the most devastating plant pathogens and represent one of the largest threats to food security (Steinberg and Gurr 2020). Efforts to combat and control fungi are constantly fraught with challenges from novel strains and/or host jumping (Fones et al. 2020). Part of the reason that fungi are such difficult pests to manage is their genomic diversity. Of particular notoriety are those strains which possess accessory chromosomes (ACs). In addition to a set of “core” chromosomes, shared by all members of a given species, many isolates carry a number of additional chromosomes. These extra chromosomes are referred to as lineage specific, conditionally dispensable, or accessory (among others), and were first described by their resemblance to B chromosomes found in animals and plants (Miao et al. 1991). These ACs have high evolutionary rates, low gene density, differential codon usage, and large repeat content (Ma et al. 2010; Coleman et al. 2009; Goodwin et al. 2011; Yang *et al*. 2020). In addition to a large number of transposable elements (TEs), they also accumulate genes beneficial to the fungi. This has been demonstrated in pathogenic fungi, where effector genes that help the fungi infect its host are often found in accessory parts of the genomes (van Dam et al. 2016; Henry et al. 2021). Due to the contrasting burden of TEs and benefits of virulence-related genes, ACs are assumed to constitute a trade-off between genomic stability and pathogenicity.

Contrary to the hypothesis that ACs are maintained because of benefits during fungal colonization/infection of plants, some ACs seem to be maintained without contributing advantages to the host, persisting even when grown on benign media where the benefits of pathogen-specific genes have little or no relevance for growth. For example, in *Fusarium oxysporum* f. sp. *lycopersici*, the ACs are lost only in every 35 000^th^ asexual spore (Vlaardingerbroek *et al*. 2016). Thus, while ACs in this species are indeed tightly linked to virulence (Ma *et al*. 2010), there appears to be little selection to lose ACs when this selective pressure is removed. In contrast, in the wheat pathogen *Zymoseptoria tritici* the loss of ACs occurs more frequently, approximately once in every 50^th^ asexual generation (Möller *et al*. 2018). However, the connection between virulence and ACs in *Z. tritici* is more complex than in *F. oxysporum*, with some ACs apparently inhibiting virulence on certain cultivars (Habig *et al*. 2017). Neither of these patterns strictly support a role of trade-offs in maintaining ACs in pathogenic fungi. As such, other mechanisms must be investigated.

Here, we hypothesize that the maintenance and spread of ACs in species of the genus *Fusarium* are linked to the presence of homologs of selfish spore killing elements that were first reported in *Podospora*, known as *Spok* genes (van der Gaag *et al*. 2000; Grognet *et al*. 2014). This hypothesis arises from compelling observations from previous work. First, large numbers of *Spok* homologs are found on ACs in *Fusarium* (Vogan et al. 2019). This includes chromosomes that exhibit whole chromosome transfer between strains of *F. oxysporum* and that have strong contributions to virulence (Ma et al. 2010; Shahi et al. 2016). Second, in addition to their role in the meiotic cycle of *P. anserina*, the *Spok* genes are expressed during vegetative growth and exhibit strong toxicity (Vogan et al. 2019). Furthermore, the *Spok* genes are capable of inducing cell death across a vast breadth of phylogenetic diversity, including other sordariales (Grognet *et al*. 2014), the model yeast *Saccharomyces cerevisiae*, and even the bacterium *Escherichia coli* (Urquhart and Gardiner 2023). This suggests that their toxic function is not limited to the sexual cycle, nor by genome or species-specific context. Therefore, the SPOK proteins could act during vegetative growth in *Fusarium* to prevent the loss of accessory chromosomes that carry them through mechanisms such as non-disjunction during mitosis.

The precise mechanism of action of the *Spok* genes has yet to be uncovered, but a number of important characteristics have been discerned. Four *Spok* homologs are known from *Podospora, Spok1* from *P. comata* and *Spok2, Spok3*, and *Spok4* form *P. anserina* (Grognet et al. 2014; Vogan et al. 2019). Despite all homologs showing a high degree of similarity, *Spok2, Spok3*, and *Spok4* show no epistatic interactions among themselves, operating independently during sexual crosses (Vogan et al. 2019). Conversely, *Spok1* provides resistance to all other homologs and can kill against *Spok2*, and *Spok3*, but not *Spok4* (Grognet *et al*. 2014; Vogan et al. 2019). The SPOK proteins appear to consist of a tripartite construction. The function of the first domain is unknown. The second domain has homology to nucleases and is responsible for the spore killing, while the third domain is a kinase and is required for resistance (Vogan et al. 2019). Site-directed mutagenesis of conserved sites has determined that the second domain (henceforth the “killing domain”) possesses a lysine in the active site that is necessary for killing, while the third domain (the “resistance domain”) has a canonical kinase active site with an aspartic acid residue at position 667 in *Spok3*. The functional separation of these two portions of the protein has been verified through generation of a truncated version that is lacking the entire resistance domain, which is lethal when expressed *in vitro* (Vogan et al. 2019). How resistance is conferred remains to be discovered, whereas the mechanism of killing appears to operate via general DNA degradation (Urquhart and Gardiner 2023). Of particular note is that this protein architecture, a nuclease domain associating with a kinase domain, is widely distributed across different domains of life (Zhang et al. 2016). It is proposed that this specific class of nuclease domain arose from bacterial transposons, later diversifying into various toxins which are involved with a range of biological conflicts (Zhang et al. 2016), including host-pathogen interactions (Van Damme et al. 2012) and gene drive systems (Lorenzen *et al*. 2008).

*Spok3* and *Spok4* are found at lower frequencies in *P. anserina* than *Spok1* or *Spok2* and are located at different genomic locations in different strains (Vogan et al. 2019). This is due to the fact that these two genes are situated within a giant transposable element, initially refereed to as the “*Spok* block”, now renamed to *Enterprise* (Vogan et al. 2019, 2021). Along with a handful of other large transposons, *Enterprise* became a founding member of the superfamily of transposons called *Starships*. These TEs average around 110 kb in size and are noteworthy for the fact that they mobilize a large number of fungal genes both within and between species (Gluck-Thaler et al. 2022; Urquhart *et al*. 2024, 2023). Gene content analysis of the *Starships* has shown that *Enterprise* is not the only element to carry *Spok* genes, with numerous other elements harbouring homologs (Gluck-Thaler *et al*. 2022). However, as no other *Spok* homologs have been functionally validated to operate as meiotic drive genes and/or toxin/antidote genes, the role that these *Starship* associated *Spok* homologs play in fungal biology remains unknown.

To evaluate our hypothesis of the SPOK protein’s role in the maintenance and spread of ACs in *Fusarium*, we assessed the diversity of *Spok* homologs (henceforth *FuSpoks*) across the genus and their association with accessory genomic regions. We then determined whether a subset of these homologs can act as toxin/antidote proteins in a *S. cerevisiae* yeast system. Together, these results allow us to propose a model for the role of *FuSpok*s in the maintenance, origin, and spread of accessory chromosomes in *Fusarium*.

## Materials and methods

### Phylogenetic analysis

We retrieved 146 high-quality RefSeq genome assemblies from 24 *Fusarium* species, selecting for assemblies that have been generated using long-read sequencing technologies (Supplementary Table S1). Genomes lacking annotations were annotated by liftover annotation using `Liftoff` (Shumate and Salzberg 2021), with the closest related reference genome with gene annotations used as the reference. This collection of *Fusarium* genome assemblies were stored in a `Mycotools` (Konkel and Slot 2023) database, which provided the framework for the identification of SPOK homologs and phylogenetic analysis. Such homologs were found using an HMM profile built from a collection of 250 protein sequences of the meiotic drive suppressor kinase protein family (IPR052396) present across *Fusarium*. The alignment of these reference amino acid sequences was created using `MAFFT` and `hmmbuild` was used to construct the final HMM model (Eddy 2011). Putative homologs were identified across this set of genomes using this HMM model and `hmmersearch`, with results based on a minimum e-value (0.001) and sequence similarity threshold (> 30%) to the query. Protein sequences were generated based on these filtered hits and which represent full protein sequences based on existing genome annotations. However, coding sequence annotations of *Spok* genes are often incorrect and contain false introns. We retrieved the nucleotide sequence for each putative *Spok* gene and identified the existing reading frame, independently of existing gene annotations. In total, we recovered 437 putative *Spok* homologs from 14 *Fusarium* species (Supplementary Table S2).

In addition to this set of *Spok* homologs, we included a set of 33 manually curated *Spok* protein sequences from *P. anserina* (n=4), *F. oxysporum* (n=13), *Fusarium poae* (n=15), and *Fusarium vanettenii* (n=1). This set of protein sequences were aligned using `MAFFT` and trimmed using `clipkit` (Steenwyk *et al*. 2020). We excluded 56 putative homologs prior the phylogenetic analysis if a homolog was i) missing catalytic core of resistance domain, ii) missing most or all of the coiled-coil domain, iii) the sequence similarity to any of the reference *Spok* sequences was too low (< 30%). Using `IQ-tree` (Minh *et al*. 2020), substitution models were compared, and a maximum-likelihood phylogeny was constructed with ultrafast bootstrap approximation. The final phylogeny was visualized using `iTOL` (Letunic and Bork 2024).

### Repeated sequencing errors of *FuSpoks*

Because of their high similarity, we did not decide on any particular *FuSpoks* from *F. poae* strain DAOMC 252244 to amplify (this was also a result of the flanking regions being too similar to design unique primers). We were able to amplify *FPOAC1_003985* placed on chromosome 2, but consistently found an indel in the amplified sequence: we found one more G (five rather than four) than the reference sequence at position 1848-1851. Looking at the nucleotide alignment to the other *FuSpoks* found in the genome, they all carry five rather than four Gs at this position. Note that the gene model on NCBI contains an intron that spans this region.

To investigate why this discrepancy existed, we downloaded the Nanopore and Illumina sequence reads for the BioProject PRJNA578270 from the SRA (SRR13483968 for the Nanopore data and SRR13023856 for the Illumina data), and aligned these to the published reference genome using Minimap2-2.24 (r1122) (Li 2018, 2021). For both sequence data sets we used samtools 1.14 (functions fixmate, sort and index with default options) (Danecek et al. 2021) followed by Picard tools MarkDuplicates 2.27.1 (Pic 2019) with options `REMOVE_DUPLICATES=true` and `CREATE_INDEX=true`. The resulting BAM files were visualized using IGV (Robinson et al. 2011; Thorvaldsdóttir et al. 2012; Robinson et al. 2017). In the Nanopore reads, the position was variably called with either four or five Gs in sequence. The Illumina reads had a drastic drop in coverage at this position, with no reads spanning the sequence of Gs. This made us draw the conclusion that the reference sequence is actually wrong, caused by a sequencing error in Nanopore. Likely, it is a combination of high GC content and repeats (Delahaye and Nicolas 2021).

The finding that the reference sequence was wrong about a this sequence made us question a second (reportedly) variable site in the *FuSpok* sequences of this genome. About half of the copies predict a frameshift caused by a single base pair deletion of a G. We repeatedly found no Illumina reads covering the region, for the copies with the reported base pair deletion. In the Nanopore data, we found that these sites were variable, with around half of the reads having single rather than double Gs. There were also an inordinate amount of sequences covering the homologous site in the full copy sequence (with two GGs) on Contig_2. We thus concluded that the premature stop codon reported for these *FuSpoks* were technical errors during sequencing.

### Fungal Strains and culturing conditions

For all experiments involving the expression of *Spok* genes in yeast, we used a modified version of the yeast strain BY4742, for which we replaced the *TRP1* gene by insertion of *URA3*. See ***Construction of yeast strain*** for details. Yeast were grown on Yeast Extract – Peptone – Dextrose (YEPD) media at 30°C before transformation.

For amplification of a *Spok* homolog from *F. vanettenii* isolate 77-13-4, the strain was acquired from the Agricultural Research Service culture collection (NRRL), under the identifier NRRL 45880. The fungus was grown on potato dextrose agar at 27 °C for 48 hours, and the spores were collected by scraping the surface of the fungal growth with a flame sterilized loop and dipping it in an Eppendorf tube with 50 µL water. The spore solution was spread uniformly on a Potato dextrose agar (PDA) plate covered with cellophane film and incubated at 27°C for three days. Mycelia was then harvested by scraping off the cellophane and stored for DNA extraction. For *P. anserina*, strain Wa28+/−was used for DNA extraction. It was grown on plates of PASM0.2 covered with cellophane. The fungal material was harvested by scraping mycelium from the surface of the cellophane and placing 80–100 mg of mycelium in 1.5 ml Eppendorf tubes, which were then stored at −20°C.

Wild strains of *P. anserina*, originally collected and maintained by the Laboratory of Genetics at Wageningen University, were used for experiments involving temperature effects (Supplementary Table S3). Sexual crossings of *P. anserina* were performed on Henks Perfect barrage medium (HPM) (Vogan *et al*. 2019) by confronting strains of opposite mating type. Cultures are maintained at 27°C under 70 % humidity and a 12:12 dark:light cycle. Sexual structures (perithecia) can be observed after approximately 10 days. Perithecia can be harvested and dissected to evaluate the occurrence of “spore killing”, or allowed to naturally shoot ascospores, which can be harvested and germinated to obtain sexual progeny. Germination is conducted on PASM2 media supplemented with 5 g/L ammonium acetate (van Diepeningen *et al*. 2008). To assay the effect of temperature on the killing action of the *Spok* genes, plates were instead incubated at range of temperatures from 22–25°C.

### Expression of *FuSpoks* in yeast

#### Genes investigated

In addition to *Spok3* from *P. anserina*, we investigated the following genes: *NECHADRAFT_82228, FOXG_14281, FOXG_14774, FOXG_07107, FuSpok1, FuSpok2*.

#### Construction of yeast strain

We replaced the *TRP1* gene in BY4742 (Baker Brachmann *et al*. 1998; Winston *et al*. 1995) by first amplifying *URA3* from plasmid DNA with primers that had 40 bp homology to the flanking regions of *TRP1* within the BY4742 genome, and 24–26 bp homology to *URA3*. The resulting PCR product was transformed into BY4742 using the lithium acetate/single-stranded carrier DNA/PEG method of transformation described by Gietz and Schiestl (2007), slightly modified to use overnight culture. Transformed culture was plated on synthetic dextrose (SD) medium lacking uracil (SD-U). Colonies were picked and streaked on SD-U and SD lacking tryptophan (−W), and one colony which grew on SD-U but on SD-W was chosen as our strain to use for the *Spok* expression assays.

#### Vector construction

We extracted DNA from *F. vanettenii* (isolate 77-13-4) using the Quick-DNA™Fungal/Bacterial Microprep Kit (ZymoResearch) and confirmed fungal DNA presence with ITS1 and ITS4 primers. DNA from *F. poae* strain DAOMC 252244 was provided by Dr. David Overy, and DNA from *F. oxysporum* f. sp. *lycopersici* strain 4287 by Dr. Antonio Di Pietro. *Spok3* from *P. anserina* was amplified from previously obtained plasmids (Vogan *et al*. 2019).

The plasmid P001, a pRS413 derivative (Mumberg *et al*. 1995), includes a 2-micron origin, AmpR (for *E. coli*), TRP1 (for *S. cerevisiae*), and a galactose-responsive *pGAL1–pCYC1* hybrid promoter (Guarente *et al*. 1982) (Supplementary File 1). The *GFP* gene in P001 was replaced by *Spok* genes in our project.

High Fidelity Phusion polymerase (ThermoFisher) was used for PCR, except for two gap repair reactions where Q5 2x mastermix was used. Primers used are found in Supplementary Table S4. Primers were designed for restriction enzyme sites to amplify genes of interest from genomic DNA, targeting flanking regions where *FuSpok* sequences diverge. PCR products and the P001 backbone were digested, ligated, and transformed into 5-*α* Competent *E. coli* C2987H (New England Biolabs) for selection. Correct plasmid construction was confirmed by colony PCR and Sanger sequencing.

Purified plasmids (QIAGEN plasmid purification midi kit) were then used as templates for gap repair PCR, designed with long primers for precise integration into the P001 backbone. The digested plasmid and amplified gene were transformed into yeast using a lithium acetate/PEG method (Gietz and Schiestl (2007)), and colonies were selected on SD-W agar plates. Exceptions to this approach was *FOXG_14774* from *F. oxysporum* f. sp. *lycopersici*, which were amplified directly from genomic DNA and used for gap repair.

#### Site-directed mutagenesis

For the majority of the genes, we mutagenized the conserved aspartic acid residue into alanine through whole plasmid PCR of the constructed plasmids. The mutagenized plasmid was then cloned in *E. coli* and used as template for gap repair as described above. For *FOXG_07107* and *FOXG_14774* we instead made two separate PCR reactions for mutagenesis, producing one product for the first part of the gene (up until and extending 21 bp beyond the mutagenized base) and one product for the second part of the gene (starting 22 bp upstream of the mutagenized site and continuing to the end of the gene). These two products were combined with the plasmid backbone for a three-segment gap repair. For details of specific site mutations, see Supplementary Table S5.

#### Growth assay

For assays of *Spok* toxicity, yeast containing the plasmids of interest were grown on SD-W for two days, after which single colonies were picked into sterile water, serially diluted, and spotted onto SD-W with either 2 % glucose or galactose. Plates where incubated at 30°C for either two days (glucose plates) or three days (galactose plates) before photos were taken. For experiments involving the effect of temperature on *Spok3* toxicity, plates were instead incubated at 22°C.

## Results and Discussion

### *FuSpoks* are rapidly diverging and enriched in accessory genomic regions

Casual observations have indicated that the homologs of the *Spok* genes in *Fusarium* are associated to the ACs and/or accessory regions of core chromosomes in some strains (Vogan *et al*. 2019). To determine if this is a general principle, and thus if there may be a causal connection between these genomic features and the *FuSpoks*, we assembled a collection of 146 high-quality genomes from across the genus. In our previous research we had observed that it is difficult to accurately assemble *Spok* gene family sequences from short-read assemblies alone. This is likely due to a combination of high similarity among homologs, gene conversion events, and a propensity to be flanked by repetitive sequences, such as TEs (Vogan et al. 2019; Ament-Velásquez *et al*. 2022). Thus, to obtain an accurate assessment of the number and diversity of *FuSpoks*, we restricted our search to only genomes produced with long-read technologies. Unfortunately, this led to another unforeseen pitfall regarding genomes produced with Oxford Nanopore Technologies (ONT). A number of *FuSpok*s were annotated as possessing introns in regions of the gene known to be critical for function, while the *Spok* genes in *Podospora* have no known introns in the coding region (Vogan et al. 2019). Manual inspection often revealed that the introns were predicted when stop codons were present in the sequence, suggesting that the genes with introns may in fact be pseudogenized. However, in some cases (particularly those involving *F. poae*), we found that while the ONT reads indicated that a stop codon was present in the gene, reads produced with Illumina technology did not. This is likely a result of repeatable errors in base calling that ONT is susceptible to (Delahaye and Nicolas 2021). As a result, we annotated protein sequences ourselves from all genomes, assuming no introns. Cases where stop codons were found prematurely were excluded from the final set, see Methods for specific details.

This approach yielded 437 *FuSpok* homologs from 95 genomes (Supplemental Table S2). The genome with the highest number of *FuSpok*s was *F. poae* strain 2516 with 25 homologs, while many other strains were found to have none (Supplementary Table S1). To assess whether *FuSpok* genes were specifically associated with ACs or were present at the edges of chromosomes (500kb), which are often enriched for accessory genomic content (Yang *et al*. 2020), we focused on a subset of 18 *Fusarium* genomes for which information on ACs was readily available. Almost a third (n=106) of the *FuSpok*s identified in this analysis (n=352) were found on ACs compared to those present on core chromosomes, small contigs, or on unidentified chromosome in the assemblies. We found that the proportion of *FuSpoks* on ACs compared to on core chromosomes was significantly higher than the proportion of non-*Spok* genes found on ACs in 11 out of the 18 *Fusarium* genomes (genes associated with the ends of contigs, small contigs, or *Starship* elements were not included) (Figure 1). While our binomial test demonstrates an association between *FuSpok* genes and ACs, it does not establish causation. We explore potential explanations for this distribution pattern in the section

**Figure 1.**
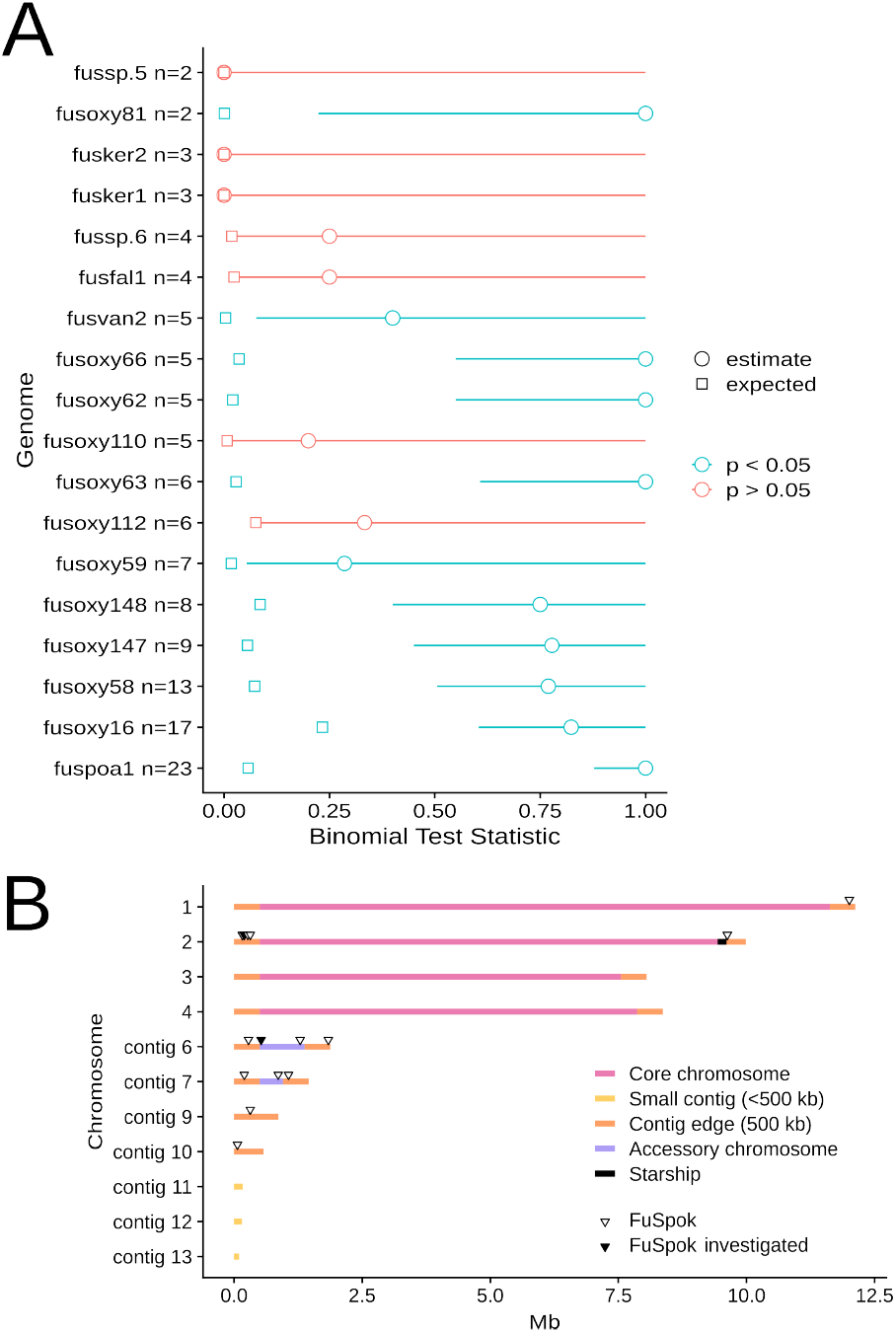
A) *FuSpoks* are enriched in the accessory regions of *Fusarium* genomes. The proportion of genes on the accessory regions of each genome is shown as squares, and the results of the binomial test for each *FuSpok* distribution is visualized with 95% confidence intervals, p-values Bonferroni corrected for multiple tests. B) Location of *FuSpok*s in *F. poae* strain DAOMC 252244. Chromosomes/contigs are coloured based on their reported identity in Witte *et al*. (2021), and small contigs or contig edges.

### On the origin of accessory chromosomes in *Fusarium*

In *P. anserina*, the three *Spok* genes are highly similar, yet have functionally diverged to act as independent toxin/antidote genes (Vogan *et al*. 2019). It is thus possible that all *FuSpok*s in a given genome represent unique toxin/antidotes. To investigate patterns of gene family diversification of the *FuSpoks*, a phylogeny of all 437 *FuSpok* homologs, as well as 29 *FuSpok* genes from our reference set was generated (Figure 2A). The topology of this phylogeny is generally consistent with existing *Spok* gene phylogenies (Grognet *et al*. 2014; Vogan *et al*. 2019), and can be separated into four main clades. *FuSpok* genes from *F. oxysporum* are prevalent across all clades, while homologs from the other species display sparser distributions, which may be due to the lower number of genomes available. One notable exception is *F. poae*, which exhibits a large number of homologs in clade IV. This is of particular interest as *F. poae* is closely related to *F. graminearum* and shares its general genome structure of having four large core chromosomes. However, unique to this species is the presence of multiple ACs (Witte *et al*. 2021), upon which many of the *FuSpoks* are located (Figure 1B). Even more striking, is the fact that we included 8 *F. graminearum* genomes in our analysis, none of which contained *FuSpok* homologs that passed our criteria (Supplementary Tables S1). This provides strong circumstantial evidence that there is a link between the origin of ACs and *FuSpoks* within a given species/lineage.

**Figure 2.**
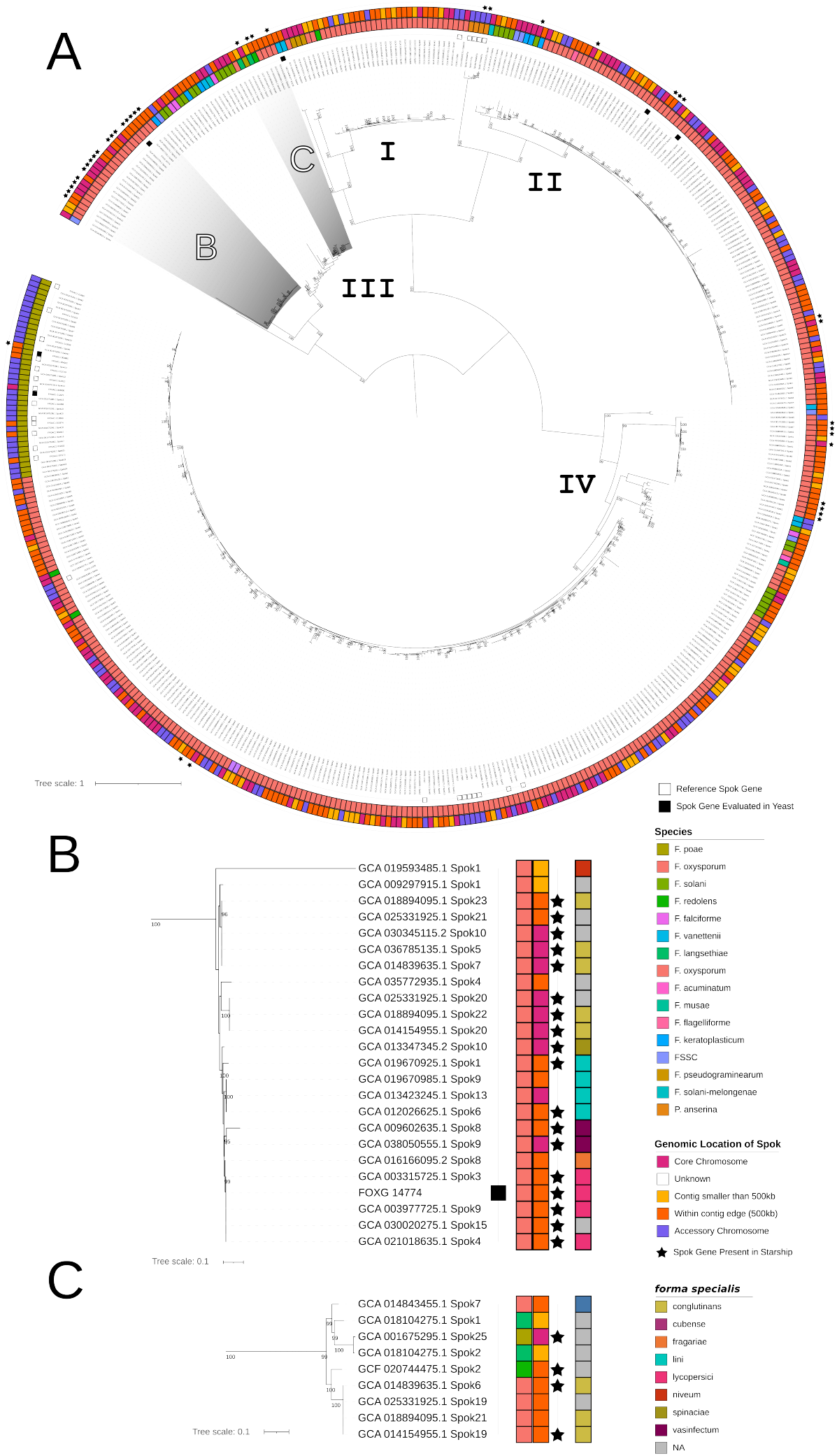
A) Maximum-likelihood phylogeny (mid-point rooted) of *FuSpok* gene homologs. The support values are ultrafast boot-strap (n=1000) approximations from `IQ-tree`, with only bootstraps with a value of >= 95 shown. Tips annotated with a black square indicate *Spok* genes that are part of the reference set (**Supplementary Table S5**), with filled squares denoting the *FuSpok* genes which were assayed in this study. The innermost coloured ring of annotations denotes species identity of the *Fusarium* genomes within the phylogeny. The outer-most coloured ring contains annotations of the genomic locations of *FuSpok* genes. Tips annotated with stars denote *FuSpoks* which were found inside of *Starship* elements. We also included a highlight for the clade containing the majority of *F. poae FuSpok* genes, which are almost exclusively present in accessory chromosomes. Certain clades enriched with *FuSpok* genes that are present within *Starships*, and are shown in sub-figures B) and C). Additional coloured annotations demonstrating the *forma specialis* designations for *F. oxysporum* genomes.

As the most well represented *Fusarium* species, most *FuSpoks* were retrieved from *F. oxysporum* genomes. Many of these *FuSpoks* were very similar in terms of sequence similarity and likely represent orthologs. As ACs are linked to the ability of specific lineages of *F. oxysporum* to infect specific plant hosts, we expected to see associations between *forma specialis* and the phylogenetic distribution of the *FuSpoks*. While there is a large amount of variation in copy number of the *FuSpoks* in specific assemblies (Supplementary Figure S1; Supplementary Table S1), no other broad trends in the *FuSpok* phylogeny of *F. oxysporum* are apparent (Supplementary Figure S2). However, more fine scaled analyses of AC synteny among the strains may be required to discern such patterns.

Given the incongruity of the *FuSpok* phylogeny with known relations among *Fusarium* species (Vogan et al. 2019), and the previous observations that the *Podospora Spok* genes are present on *Starships* (Vogan et al. 2021), we explored the association of these massive transposons with the *FuSpoks*. A total of 39 of the 437 *FuSpoks* in our phylogeny were found to be within known *Starships* (Figure 2). The majority of the *FuSpoks* within *Starships* were found within *F. oxysporum* genomes. Specific clades of *FuSpok* genes were enriched within *Starship* regions (Figure 2B), with *Spok* genes from different *Starship* families present in different *Fusarium* species (Figure 2C). Interestingly, one *F. poae FuSpok* homolog, that is present within a *Starship*, is found within clade IV,(Figure 2A). As the majority of other *FuSpoks* in this clade are from *F. oxysporum*, it suggests that the *FuSpoks* may have been introduced to *F. poae* by a *Starship* from *F. oxysporum* and subsequently underwent a broad expansion. However, as the association between *FuSpoks* and *Starships* is somewhat transient (Figure 2), we cannot rule out the possibility that this *FuSpok* was recently acquired by a *Starship* instead.

As the identification of *Starships* can be complicated by a number of factors (Gluck-Thaler and Vogan 2024), our annotations of these elements is likely incomplete. Nevertheless, our phylogeny of *Spok* homologs suggests that *FuSpoks* are rapidly gained and lost by *Starships* and that the horizontal transfer of *Starships* does influence *FuSpok* distributions. For example, the single *F. poae FuSpok* in clade III may have been transferred from *Fusarium langsethiae*. Given that the horizontal transfer of *Star-ships* has been linked to the outbreak of novel fungal diseases in crops (Bucknell *et al*. 2024), whether the presence of *FuSpoks* on *Starships* influences their transmission requires further attention.

### *Spok3* from *P. anserina* operates as a toxin and antidote in *S. cerevisiae*

Previously, we investigated the function of the *Spok* genes through genetic manipulations within *P. anserina*. While this provided numerous key insights to the functionality of the *Spok* genes, it had a number of drawbacks. In particular it was difficult to assay the toxin function of the genes as it was discovered that the *Spoks* not only kill developing sexual spores, but are active vegetatively. Thus, investigations of point mutants on the resistance function of the genes had to be inferred from a lack of transformants, rather than through observations of spore killing directly (Vogan *et al*. 2019). To remedy this issue, we developed a killing assay using the model yeast *S. cerevisiae*.

We utilised a vector based approach wherein the full length *Spok3* gene was placed under inducible expression via the Gal1 promoter. As compared to a control vector expressing GFP, a minor reduction in growth was observed, suggesting that the full length *Spok3* gene has a slight deleterious effect on yeast growth (Figure 3). In *P. anserina*, a D677A point mutation, which is located within the active site of the resistance domain (Vogan *et al*. 2019), was shown to reduce the resistance function of the protein. Congruently, this same point mutation reduced yeast growth substantially (Figure 3). This result indicates that the *Spok3* gene is an active toxin and antidote within *S. cerevisiae*. This agrees with previous results by Urquhart and Gardiner (2023) who obtained similar results with assays performed with the *Spok1* gene from *P. comata*. We confirmed that two additional modifications of *Spok3* operate similarly in the yeast assay as in *P. anserina*, namely a truncated version of *Spok3*, which has a stop codon inserted after amino acid 490, and the K240A point mutation in this truncated version. The truncation removes the entire resistance domain, while the point mutation is in the active site of the killing domain and disables the toxicity function of the gene (Supplementary Figure S3). Strikingly, this truncated version of *Spok3* inhibited yeast growth to a much stronger extent than the D667A point mutation, suggesting that some resistance activity is still maintained in the mutated active site.

**Figure 3.**
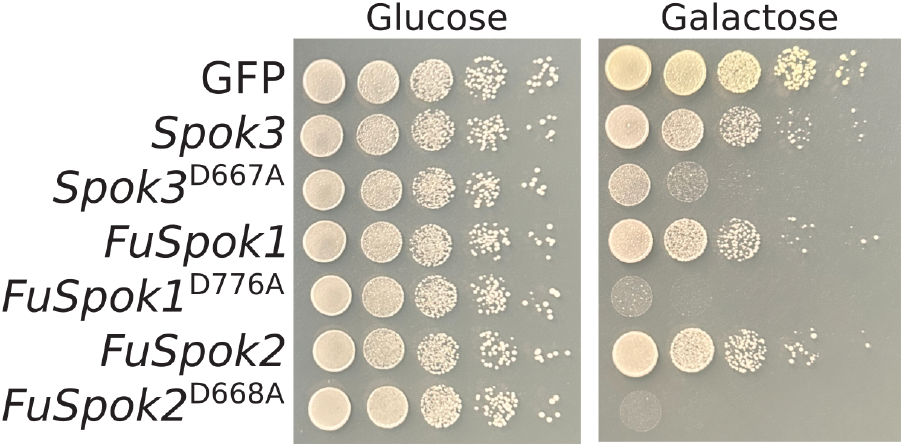
Heterologous expression of *Spok* genes in *S. cerevisiae*.

### *FuSpok* genes function as toxin/antidote proteins

Although the *Spok* homologs distributed throughout *Fusarium* show a high degree of similarity to the *Podospora* genes, including conservation of characterized catalytic amino acids (Supplementary File 2), as yet it is unknown whether they possess toxin and/or antidote function. To address this, we selected six homologs from three different species to investigate with the yeast assay (Table 1). We chose genes based on a combination of factors, including phylogentic diversity and genomic location. Additionally, we selected the *NECHADRAFT_82228* (accession: XM_003041460.1) gene from *F. vanettenii* as it was investigated previously and shown to possess toxin activity, but no antidote activity (Grognet et al. 2014).

**Table 1.** *FuSpoks* which toxicity we evaluated in yeast. The **Result in yeast** column refers to the effect of the protein with a point mutation leading to the replacement of a predicted aspartic acid residue with an alanine (corresponding to the D667A mutation previously investigated for SPOK3).

| Species | Strain | Locus Tag | Gene Name | Result in yeast | GenBank accession |
| --- | --- | --- | --- | --- | --- |
| *Fusarium vanettenii* | NRRL 45880 | *NECHADRAFT_82228<sup>a</sup>* | Unnamed | No effect | NA |
| *Fusarium oxysporum* f. sp. *lycopersici* | Fol4287 | *FOXG_14281* | Unnamed | No effect | NA |
| *Fusarium oxysporum* f. sp. *lycopersici* | Fol4287 | *FOXG_14774* | Unnamed | No effect | NA |
| *Fusarium oxysporum* f. sp. *lycopersici* | Fol4287 | *FOXG_07107* | Unnamed | No effect | NA |
| *Fusarium poae* | DAOMC 252244 | *FPOAC1_003985* | *FuSpok1* | Toxicity | PQ540985 |
| *Fusarium poae* | DAOMC 252244 | *FPOAC1_012971* | *FuSpok2* | Toxicity | PQ540984 |

<sup>a</sup> NCBI gene model includes putative false intron

We selected two genes from the species *F. poae*, one located on a core chromosomes (*FPOAC1_003985* on chromosome 2), and the other on an accessory contig (*FPOAC1_012971* on contig 1). These genes were of interest for two primary reasons. First, there are a large number of *FuSpoks* in this strain, many of which are highly similar, suggesting recent expansion. Second, the presence of ACs is unique in this species among close relatives. When copies of the *FuSpok*s that possess point mutations in the resistance domain were introduced to *S. cerevisiae*, a strong inhibition of growth was observed, greater even than what was obtained with *Spok3* (Figure 3). Given their function, we name these genes as *FuSpok1* for *FPOAC1_003985* and *FuSpok2* for *FPOAC1_012971*, see Table 1. This result provides strong support for a role of *FuSpok*s in the maintenance and/or spread of accessory chromosomes in *Fusarium*. In conjunction with the results from the phylogeny, we can infer a scenario where the acquisition of a *FuSpok* gene from *F. oxysporum* provided the means for the evolution of ACs in *F. poae*. Additionally, given the large number of highly similar homologs in clade IV from other *Fusarium* species, it suggests that hundreds of *FuSpok*s may be functional toxin/antidote genes (Figure 2). Thus supporting our main hypothesis that the *FuSpok*s are key players in the evolution of ACs across the genus.

Confoundingly, results from the other four investigated genes (Table 1) showed no impact on yeast growth, including *NECHADRAFT_82228* (Supplementary Figure S4). A number of factors may be responsible for these observations. First, despite conservation of known active sites, these genes may no longer function due to changes elsewhere in the proteins. This is partially supported by the fact that *NECHADRAFT_82228* displayed no resistance function in previous experiments, despite appearing to have a fully functional domain (Grognet et al. 2014). However, the fact that we did not observe inhibition in yeast growth, even of the unmutated copy, while previous experiments demonstrated toxic properties in *P. anserina*, may indicate that our assay is not sensitive enough to capture the whole spectrum of toxin effects. We see variation in the extent of yeast growth inhibition between *Spok3* and the *F. poae* homologs, implying that not all proteins impact *S. cerevisiae* to the same degree. It is therefore possible that the NECHADRAFT_82228 is a much weaker toxin, and that its effect is not observable under the assayed conditions. Urquhart and Gardiner (2023) observed increased sensitivity to *Spok1* when a DNA repair mutant was used, which could be a viable option for future assays of *FuSpok* genes. Additionally, as the truncated *Spok3* exhibited a much stronger toxic effect than the point mutation, it is possible that mutating the aspartic acid residue has minimal impact on resistance function for some of the homologs. Lastly, proteins may not be expressed or produced correctly from the constructed vectors for unknown reasons. Alternatively, it may be the case that the *FuSpok*s do not generally operate as toxin/antidote genes, but that their native function has been co-opted for selfish behaviour separately in both *Podospora* and *F. poae*. However, we find this explanation unlikely given the degree of gene family expansion observed in strains like *F. oxysporum* f. sp. *lycopersici* 4287. Until an alternative function of *Spok* homologs is observed, we believe that all copies should be expected to be either toxin/antidote genes or non-functional pseudogenes.

### Temperature effects spore killing in *P. anserina*

One of the greater mysteries surrounding the *Spok* genes is how a single protein is capable of acting as both a toxin and an antidote. In many other systems the two roles are encoded either by separate, linked genes, or by alternate isoforms of the same gene (Bravo Núñez *et al*. 2018). Although the mechanism is unknown, it follows that there should be two separate forms of the SPOK protein, one responsible for toxicity, and the other for the antidote function. In systems where separate isoforms of the same gene encode the two functions, the antidote isoform is often shorter-lived than the toxin. For example, this is how spore killing is enacted in the model yeast *Schizosaccharomyces pombe* (Nuckolls *et al*. 2017; Hu *et al*. 2017). This is necessary in order to create an asymmetry whereby cells with the toxin/antidote system are protected from self-toxicity, while those cells that do not inherit the gene(s) are killed. Previous work by van der Gaag (2005) supports this model of asymmetric stability for the *Spok* genes in in *P. anserina*.

Observations of spore killing between numerous strains indicated that the effect is inhibited at low temperature in some crosses (van der Gaag 2005). Together with more recent results, which connected observations of spore killing among strains with the specific presence and location of the *Spok* genes (Vogan *et al*. 2019), this observation suggests that either *Spok2, Spok3*, or both lose the killing function at 22°C. To investigate this phenomenon, we performed crosses of various wild type strains of *Podospora* at 22°C, designed to isolate the effect of single *Spok* genes. Although not all combinations of *Spok* genes are present in wild strains, specific combinations result in spore killing due to only one of the homologs. For example, a cross between Wa63 (possessing *Spok2*) and Wa28 (possessing *Spok2* and *Spok3*) results in spore killing driven by *Spok3* gene alone (Vogan *et al*. 2019). Results from these crosses indicate that both *Spok2* and *Spok3* lose their killing function below 25°C, while *Spok1* and *Spok4* maintain killing function down to at least 22°C (Supplementary Table S6). To verify that the *Spok* genes were still functional in these crosses and that the loss of killing was not due to other effects, such as spontaneous mutations, progeny were isolated from crosses conducted at 22°C and evaluated for their ability to induce spore killing at standard conditions of 27°C. These progeny exhibit the expected spore killing phenotypes, indicating that it is indeed the low temperature that inhibits the killing function (Supplementary Table S6).

The loss of killing could be explained by one of two alternative mechanisms. Either low temperature inhibits the toxic function of the SPOK protein, or alternatively, it could stabilize the antidote function such that there is no asymmetry in the degradation rates of the toxin and antidote functions. To ascertain whether the SPOK3 protein is able to operate as a toxin at low temperature, we conducted the same yeast killing assay at 22°C, instead of at 30°C as done previously. Intriguingly, the mutated version of *Spok3* showed no impact on yeast growth inhibition at 22°C compared to 30°C(Supplementary Figure S5). This suggests that toxicity is unaffected by low temperature.

These results support the model whereby cell death is induced because the antidote function decays faster than the toxin function. In *Podospora*, during the initial stages of spore formation, all daughter nuclei from meiosis share a common cytoplasm. It is shortly after spore wall formation, at which point the nuclei become isolated from one another, where sensitive spores degrade during the spore killing process (Grognet *et al*. 2014). This is consistent with a mechanism whereby the nuclei that inherit the *Spok* gene(s) produce products with toxin and antidote functions, but where antidote function breaks down faster than toxin function. Immediately after spore wall formation, the cytoplasm of all spores is infused with both toxin and antidote. However, as the antidote function quickly degrades, in spores that did not inherit the *Spok* genes, and thus cannot produce their own antidote, the toxicity induces the abortion of the spores. The effect of decreased temperature on *Spok2* and *Spok3* suggests that this condition stabilizes the antidote function for long enough that the spores survive before both toxin and antidote function eventually degrade. The fact that no effect on yeast growth inhibition was observed at low temperature also supports this model. As inhibition is only observed with the mutated *Spok* gene, stabilizing the antidote function will still not result in resistance to the toxin.

### Model for the maintenance of accessory chromosomes by *FuSpok* genes in *Fusarium*

Given the different points of evidence presented herein, we can now propose a model for the role of the *FuSpok* genes in *Fusarium*. Transcription evidence indicates that the *Spok* genes are consistently expressed, including in vegetative growth, suggesting that the entire mycelia is infused with SPOK protein (Vogan et al. 2019). To avoid the toxic effects of the protein, a nucleus must either possess a copy of the specific *Spok* gene or share a cell with a nucleus that does (Grognet et al. 2014; Vogan et al. 2019). Furthermore, the model for action of the *Spok* genes indicates that the antidote function degrades faster than the toxin function. As such, if a nucleus were to lose an AC during growth— either through non-disjunction at mitosis or other means—and become isolated in its own cell, it will lack the antidote function. Since the nucleus would have been exposed to the *Spok* product during growth, as the antidote function degrades the nucleus will experience the toxic effects, *i*.*e*. DNA damage, and break down (Figure 4). This resembles the well-described type II toxin-antitoxin systems in bacteria where a stable toxic protein is being neutralized by a more unstable antitoxin that requires constant production. The toxin-antitoxin systems in bacteria not only aid in the maintenance of the plasmid, but also of genomic pathogenicity islands, integrons, or entire chromosomes (Qiu *et al*. 2022).

**Figure 4.**
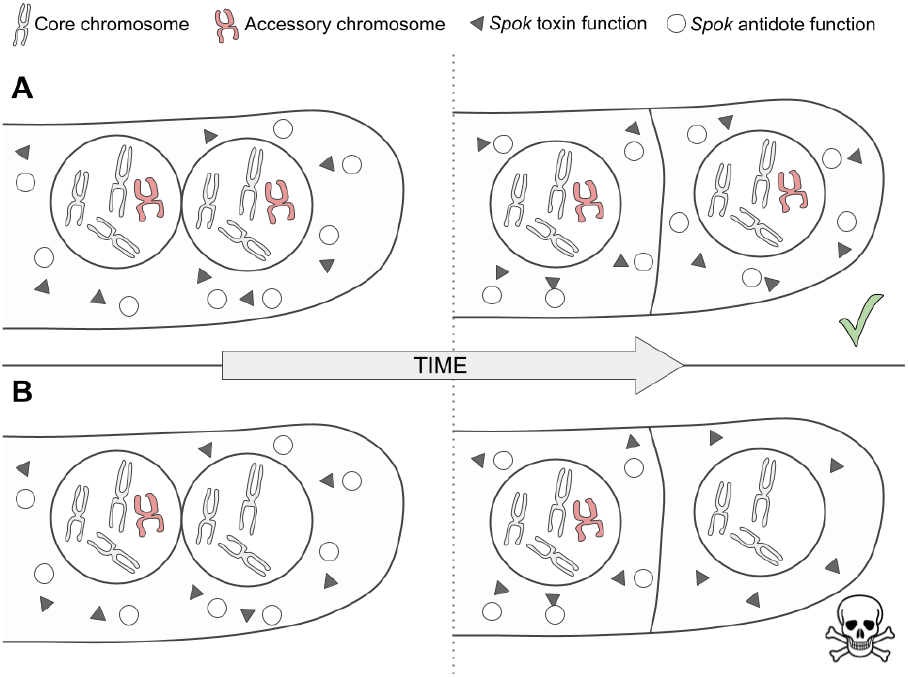
Model for the maintenance of accessory chromosomes by *FuSpoks*. Triangles represent the toxin function of the proteins, and circles represent the antidote function. Core chromosomes are depicted in grey, with ACs in red. A) A dividing cell within a hyphal tip, expressing a SPOK protein from an AC. B) A dividing cell within a hyphal tip, expressing a SPOK protein from an AC. The AC is lost during mitotic division, resulting in a cell containing *Spok* toxin, but with no ability to produce *Spok* antidote. This cell is expected to die.

So far, the understanding has been that SPOK proteins in *Podospora* only influenced transmission during meiosis, but our updated model suggests that they act throughout the entire life cycle to bias transmission. This aspect is especially relevant when considering genomic reductions after parasexual fusion, which is a likely cause of observed horizontal chromosome transfer in *F. oxysporum* (Vlaardingerbroek *et al*. 2016). In *Fusarium*, this mitotic drive has significant implications, as *FuSpoks* are located on accessory chromosomes that carry genes that increase pathogenicity of the fungus. Understanding that these chromosomes are not only selected for during host infection, but are also actively transmitted and propagated within populations provides valuable insights for predicting the spread of these pathogen-causing elements.

### On the origin of accessory chromosomes in *Fusarium*

ACs are common across fungi, but their origins remain a mystery. The presence of *Spok* genes may help solve this riddle. When a novel chromosome is formed, possibly via chromosomal fission or other structural rearrangements, in a genomic region containing a *Spok* gene, the selfish properties of the *Spok* ensure that this new chromosome cannot be lost. Alternatively, since *Spok* genes are frequently found on *Starship* transposons (Gluck-Thaler *et al*. 2022), a *Spok* gene could be brought to a novel chromosome soon after formation via transposition of a *Starship*, thus precluding the need for *Spok* genes in the background or core genome of the fungus. As *Starships* often degrade, sometimes rapidly (Urquhart *et al*. 2024), this process can be difficult to observe, but the fact that at least some *Spok* genes studied here are on *Starships* supports such a role. Selection may then lead to the movement and/or *de novo* creation of genes with roles in rapidly evolving traits, like virulence, on said chromosome. If an AC becomes beneficial due to these genes, the *Spok* genes may no longer be needed to maintain it, and could be lost or transferred to other genomic regions. Thus, while we find *Spok* genes to be enriched on ACs, this would explain why they are not a strict requirement for their existence. Other selfish properties of ACs, such as the meiotic drive observed in *Z. tritici* (Fouché et al. 2018), may also play a role in the existence of ACs that do not carry *Spok* genes.

Homologs of *Spok* genes are not restricted to *Fusarium* and *Podospora*, but are found distributed throughout a broad range of species across the Pezizomycotina. Considering *Starships* can mobilize both within and between species (Urquhart et al. 2024, 2023), their movement likely contributes to the incongruous phylogenetic distribution of *Spok* genes (Vogan et al. 2019). It remains to be answered whether there is a deep evolutionary link between the *Starships* and *Spok* genes or if *Starships* recurrently abduct *Spoks*. In either case, as the presence of *Spok* genes does not coincide directly with the presence of ACs, other factors must be in play. Nevertheless, understanding that *Spok* genes play a key role in AC evolution is an important step in determining why ACs appear in some species and not others.

## Conclusions

We have provided evidence that the role of *Spok* homologs in *Fusarium* is, at least in part, to maintain ACs during vegetative growth. Due to the large number of *FuSpoks* in given strains of *Fusarium* and the fact that there can be epistatic interactions between different homologs (Grognet et al. 2014), it is difficult to predict whether every *FuSpok* plays an active role in accessory genome maintenance. The *Spok* genes could also play a role in the parasexual cycle, which is key for the process of whole chromosome transfer in *Fusarium*, and likely plays a role in the evolution of virulence (Shahi et al. 2016). Understanding if different *FuSpok*s in a given genome have epistatic interactions and thus dictate which ACs can be transferred between strains should be a priority of future work. Although, the large number and diversity of these genes makes this a formidable challenge. Additionally, follow up work to demonstrate the role of *FuSpoks* within *Fusarium* itself is required to verify the model proposed here. This work is an important first step in deciphering the role of selfish genomic elements in the origin, maintenance, and spread of ACs and thus of virulence in devastating plant pathogens with far reaching implications to managing these important fungi for agriculture and society more broadly.

## Supporting information

Supplementary Figures and captions for supplementary tables and files

## Data availability

Strains and plasmids are available upon request. Supplementary Table S4 contains primers used.

## Acknowledgments

Dr David Overy and Thomas Witte from Agriculture and Agri-Food Canada provided DNA and sequence data from *F. poae* strain Fp157. Dr Antonio Di Pietro from University of Córdoba, Spain, provided DNA from *Fusarium oxysporum* f. sp. *lycopersici* strain 4287. Thanks to Armin Rassooli Tilehnovi for important work on exploring *Fusarium Spok* homolog diveristy.

