## Supplementary Figures and captions for supplementary tables and files for "The role of toxin/antidote genes in the maintenance and evolution of accessory chromosomes in *Fusarium*"

**S1** Distribution of Spok genes across *Fusarium* species and *F. oxysporum forma specialis* . . . . 2

**S2** Phylogeny of FuSpok genes from *Fusarium oxysporum* . . . . . . . . . . . . . . . . . . . . . . 3

**S3** *Spok3* killing assay using *S. cerevisiae* . . . . . . . . . . . . . . . . . . . . . . . . . . . . 4

**S4** *FuSpok* killing assay. . . . . . . . . . . . . . . . . . . . . . . . . . . . . . . . . . . . . . 5

**S5** *Spok* killing assay at low temperature. . . . . . . . . . . . . . . . . . . . . . . . . . . . . 6

**Supplementary table captions** **7**

**Supplementary file captions** **7**

### Supplementary Figures

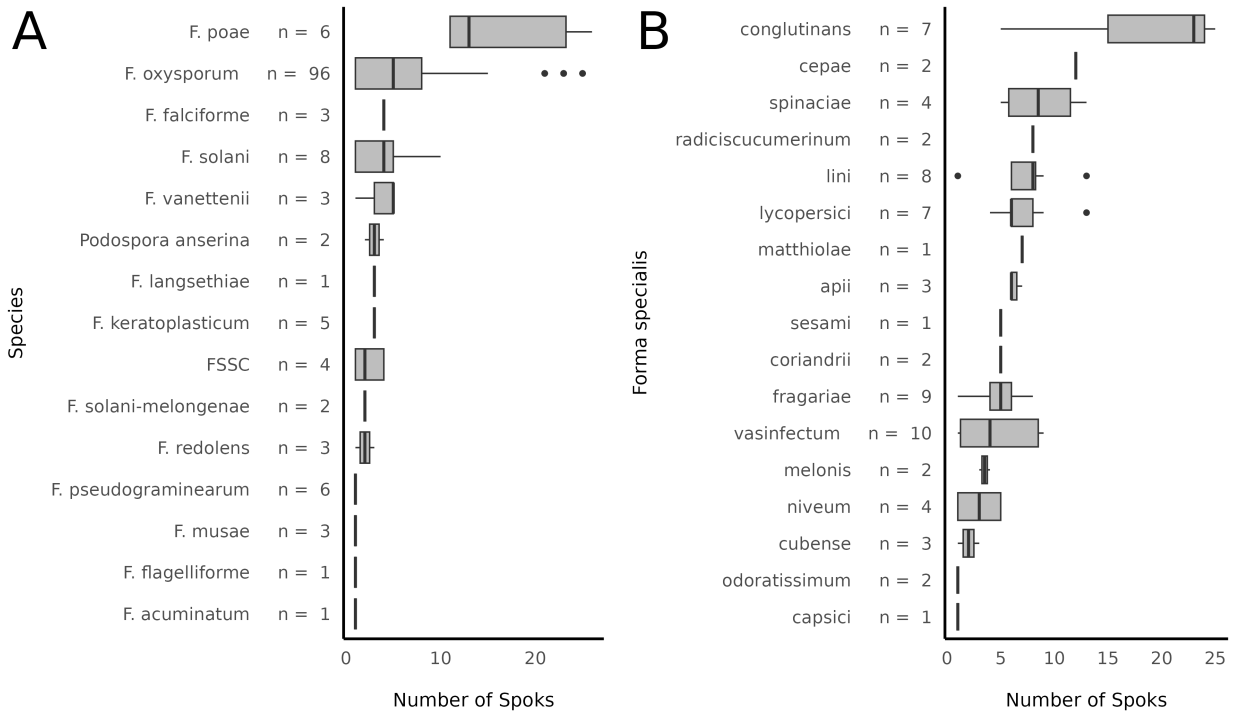


**Supplementary Figure S1**: The number of Spok genes found across (A) *Fusarium* species and different *F. oxysporum forma specialis*. The number of genomes that were considered in the search for *Spok* genes is included beside the species/*forma specialis* name.

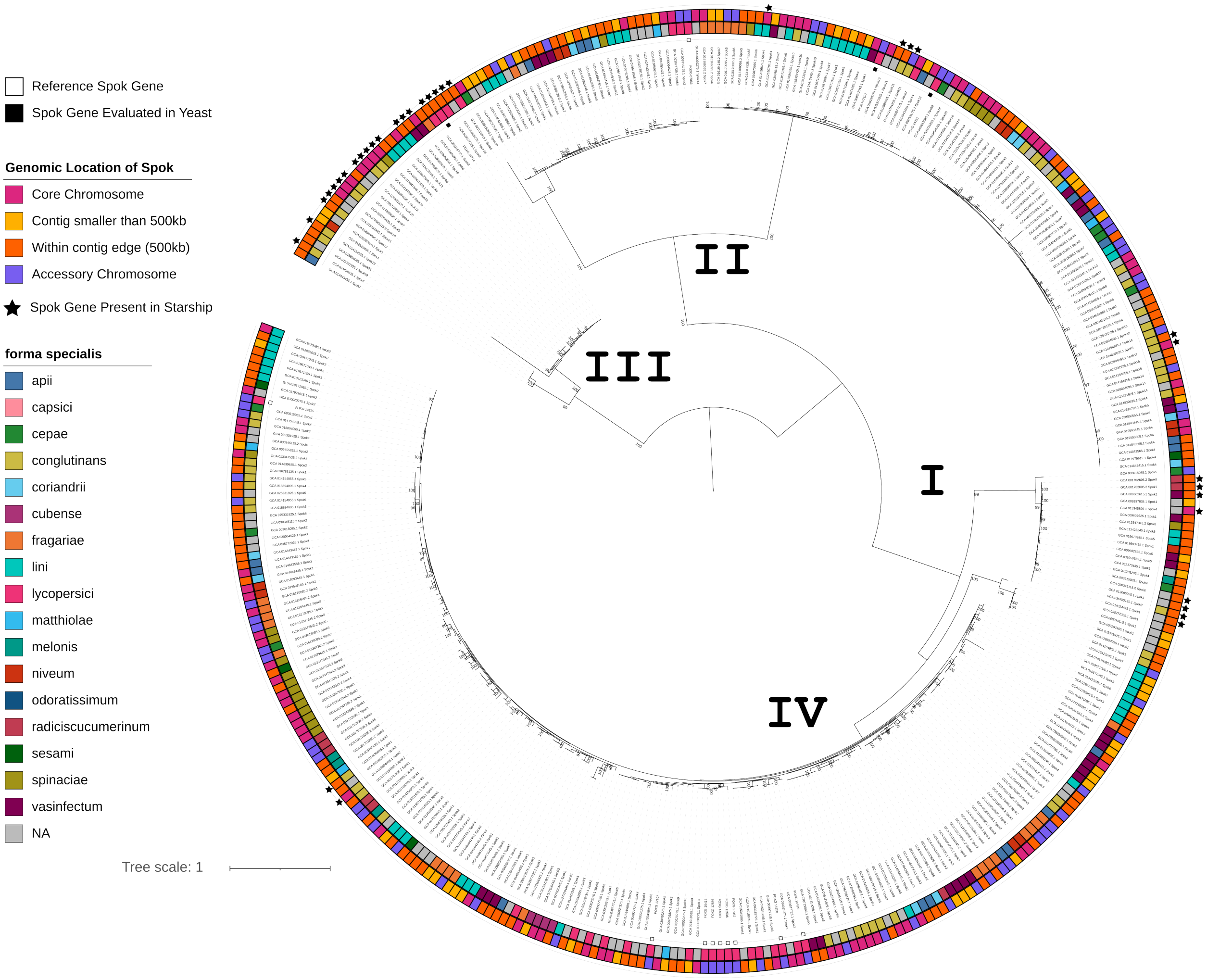


**Supplementary Figure S2**: Maximum-liklihood phylogeny (mid-point rooted) of *Spok* gene homologs from *Fusarium oxysporum*. The support values are ultrafast bootstrap (n=1000) approximations from `IQ-tree`, with only bootstraps with a value of $>= 95$ shown. Tips annotated with a black square indicate *FuSpok* genes that are part of the reference set (**Supplementary Table S5**), with filled squares denoting the *FuSpok* genes which were assayed in this study. The inner coloured ring contains annotations of the genomic locations of *FuSpok* genes, while the outer coloured ring depicts the *forma specialis* designations for *F. oxysporum* genomes. In addition, we included annotations with stars to denote *FuSpoks* which were found inside of *Starship* elements.

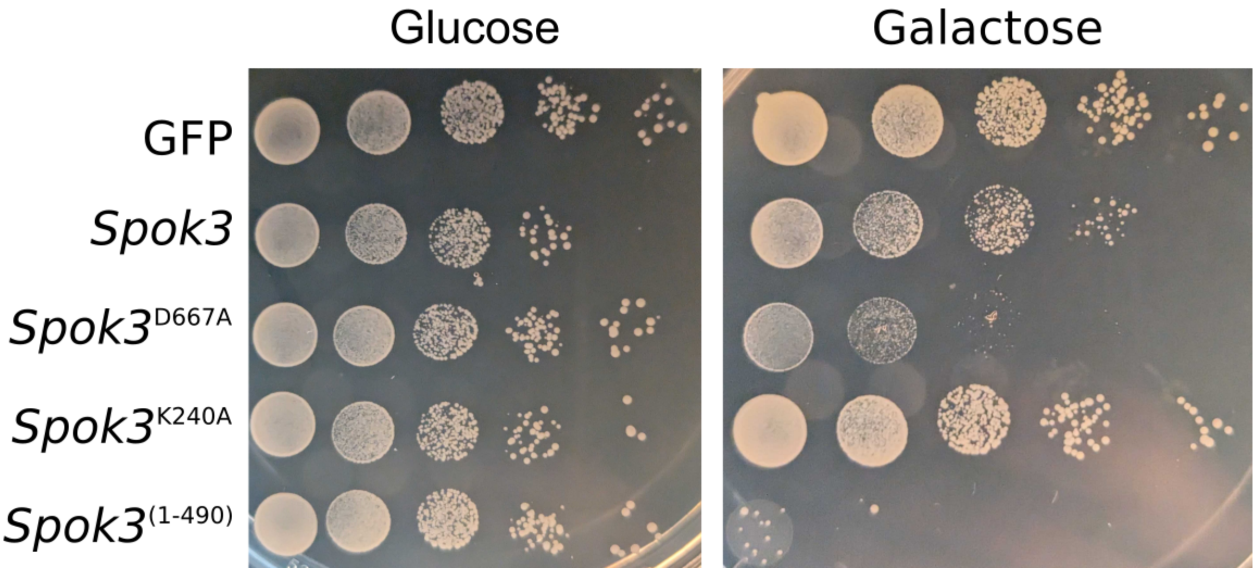


**Supplementary Figure S3**: Killing assay with wild type and mutated versons of the *P. anserina Spok3* gene expressed in *S. cerevisiae*

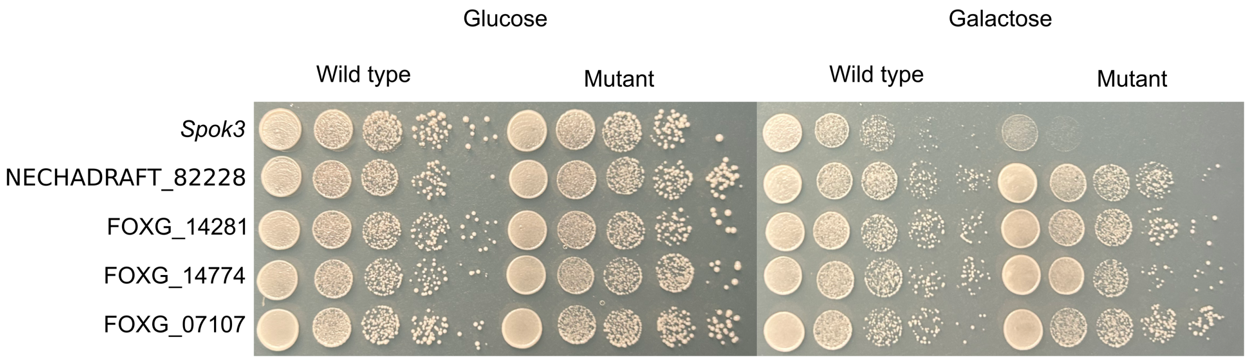


**Supplementary Figure S4**: Killing assay of various *FuSpok* genes that showed no phenotypic effects when expressed in *S. cerevisiae*. Mutant refers to strains with vectors containing *FuSpok* homologs with a D to A mutation in the active site of the resistance domain. See table S5 for details.

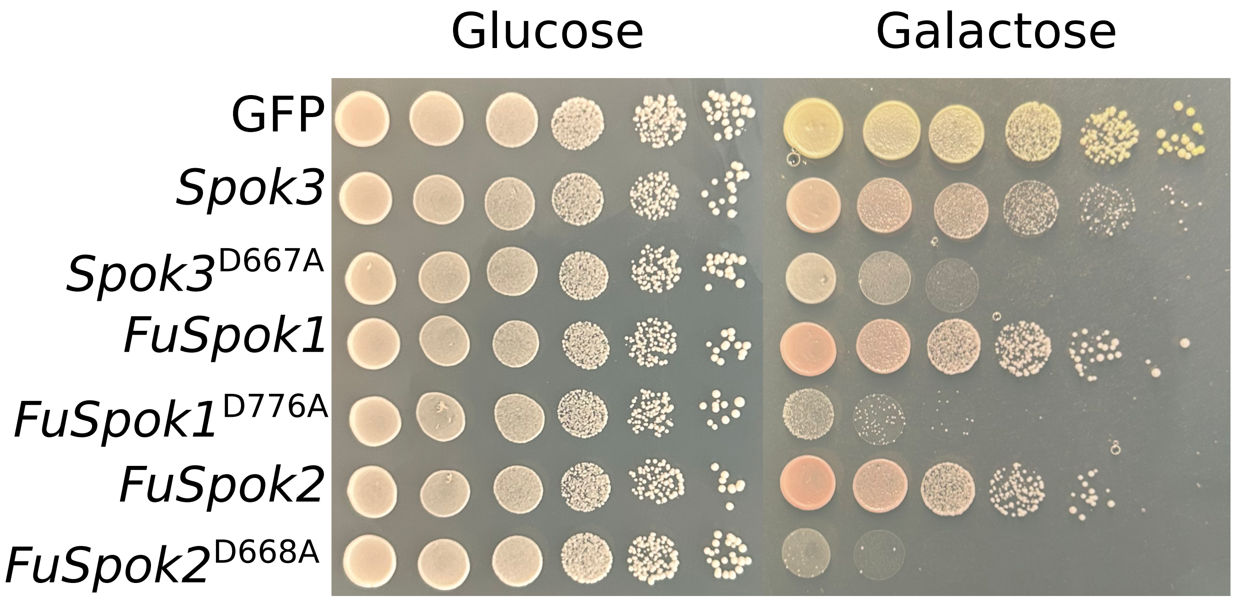


**Supplementary Figure S5**: Killing assay of various *Spok* genes conducted at 22℃. No significant difference are apparent versus growth at 30℃, except for the red colouration. This is presumably due to a lack of adenine as a result of the DNA damage induced by the *Spok* genes.

### Supplementary table captions

**Supplementary Table S1**: We identified *Spok* homologs from 145 *Fusarium* and one *Podospora anserina* genome assemblies collected from NCBI. Each genome assembly was assigned a unique genome code (the ”ome” codes from the mycotools database), generated using the first 3 letters of genus and species names, plus a numerical identifier.

**Supplementary Table S2**: The final set of *Spok* homologs and their genomic coordinates, identified within the genomes listed in Table S1.

**Supplementary Table S3**: The wild-type strains of *Podospora anserina* used for experiments involving temperature effects on spore-killing action. Originally collected and maintained by the Laboratory of Genetics at Wageningen University.

**Supplementary Table S4**: Primer sequences for amplification of *FuSpok* genes.

**Supplementary Table S5**: Details of *Spok* homolosg investigated in this study, including specific site mutations from site-directed mutagenesis experiment conducted on *FuSpoks* in yeast.

**Supplementary Table S6**: Effect of temperature on spore killing in *Podospora*. 2-spored asci indicate the presence of spore killing. Expected rates of spore killing for each cross can be found in Vogan et al. 2019. Strains 63x28_8c, 47x28_bm, 47x28_d, and 47x28_e refers to a progeny isolated from a 5-spored asci of crosses between the indicated parent strains, where 47x28_bm is from a monokaryotic spore.

### Supplementary file captions

**Supplementary file S1**: Fasta file of entire plasmid P001 sequence.

**Supplementary file S2**: Multiple sequence alignment of all *Spok* homologs. Used for generating phylogenetic trees.
